# Three-dimensional quantitative characterization of *Coxiella burnetii* infection using focused ion beam-scanning electron microscopy

**DOI:** 10.1101/2025.01.03.631259

**Authors:** Jack M. Botting, Samuel Steiner, Morven Graham, Xinran Liu, Craig R. Roy, Jun Liu

## Abstract

*Coxiella burnetii* is a highly virulent intracellular pathogen that causes acute and chronic Q fever in humans. The bacterium utilizes a Dot/Icm type IV secretion system (T4SS) to translocate over 100 effectors into host cells to facilitate biogenesis of *Coxiella*-containing vacuoles (CCVs), which are specialized lysosome-derived organelles that support bacterial replication. During replication in CCVs, *C. burnetii* undergoes a unique biphasic developmental cycle defined by two distinct cellular forms: the infectious and metabolically dormant small cell variant (SCV) that converts to a replicative large cell variant (LCV). This developmental cycle was believed to be intimately coupled with biogenesis of CCVs, but the underlying mechanisms remained unclear. Here, we combine advanced focused ion beam-scanning electron microscopy (FIB-SEM) with machine learning-based data analyses to visualize host cells infected by *C. burnetii*. We reveal significant pleiomorphism of both the bacteria and CCVs in three dimensions. We show that this technology can be leveraged to characterize CCV biogenesis defects displayed by *C. burnetii* mutants that are unable to generate the large CCV displayed by wild-type *C. burnetii*. Analysis of HeLa cells infected with a *cig2*::Tn mutant confirmed that this mutant creates vacuoles that have a defect in homotypic fusion but that the proportions of SCV to LCV in vacuoles in which this mutant resides are nearly identical to CCVs containing wild-type *C. burnetii*, which indicates the vacuole biogenesis defect displayed by the *cig2*::Tn mutant did not impact the developmental cycle intracellularly. Collectively, this study provides unprecedented three-dimensional images of the complex intracellular lifestyle of *C. burnetii*. This imaging technology also provides unique insights into the biphasic developmental cycle and will be a powerful approach to dissect CCV biogenesis defects displayed by mutant *C. burnetii*.

## INTRODUCTION

The gram-negative pathogen *Coxiella burnetii* is responsible for Q fever in humans, a zoonosis transmitted from livestock that presents flu-like symptoms and can result in endocarditis and persistent infection (1). Aerosol infection by *C. burnetii* can result from an extremely low dose of ∼10 cells and is facilitated by the bacteria being very stable in the environment (2–4). After inhalation *C. burnetii* infects alveolar macrophages, where it replicates inside a modified phagolysosome called the *Coxiella*-containing vacuole (CCV) (5). Although *C. burnetii* is an obligate intracellular pathogen it is possible to cultivate the bacteria axenically in liquid medium and on agar plates, which opens new avenues to investigate mechanisms of pathogenicity (6).

The environmental stability and infectivity of *C. burnetii* is attributed to its unusual biphasic developmental cycle (4). *C. burnetii* adopts two distinct morphologies: a dormant small cell variant (SCV) and a replicative large cell variant (LCV). Both variants are straight rods, but the SCV is much shorter than the LCV (0.2-0.5 μm versus 1 μm, respectively) and has much denser genetic material (4, 7). The cell envelope of the SCV is also decorated with extra membrane folds extending into the cytoplasm. These structures are thought to act as a form of membrane storage that facilitates the transition from SCV to LCV (7). The SCV is the infectious form that differentiates into the LCV once the bacteria establish a niche inside the host cell that is permissive for bacterial replication. Specifically, SCV-to-LCV differentiation is triggered by acidification of the CCV as it matures along the endocytic pathway that promotes fusion with lysosomes. This also serves as a signal for activation of the *C. burnetii* Dot/Icm type IV secretion system (T4SS) (5, 7–12).

The major role of the *C. burnetii* Dot/Icm T4SS and its translocated effector proteins is to remodel the CCV into a compartment allowing bacterial replication. For instance, the effectors Cig2 and Cig57 have been found to play a role important for creating a CCV that supports robust *C. burnetii* replication and the maintenance of the CCV in an autolysosomal state by promoting the fusion of autophagosomes with the CCV (13–18). T4SS function is essential for intracellular replication and CCV biogenesis (8, 14, 19, 20). CCV fusion with autophagosomes and constitutive, homotypic fusion of early-stage phagocytic vesicles containing *C. burnetii* are driven by host proteins, including SNAREs and autophagy proteins (14, 15, 21, 22). Three-dimensional imaging techniques capable of revealing specific *C. burnetii*-host interactions during infection would therefore be of great utility to understanding the mechanistic contribution of bacterial effectors and host proteins during CCV biogenesis.

Among many modalities of electron microscopy (EM) that probe pathogen-host interactions at the nanometer scale (Weiner & Enninga 2019), focused ion beam-scanning electron microscopy (FIB-SEM) is a relatively new imaging technique with significant advantages, such as high isotropic resolution (< 10 nm in x, y, and z) and fully automated operation. Importantly, these advantages are critical for subsequent data analysis to better interpret the acquired 3D data. In this study, we use FIB-SEM to image *C. burnetii* populations within host cells. We have developed machine learning-guided data analyses that efficiently characterize bacteria, CCVs, and features of host cells, which together provide extensive quantitative information about the course of *C. burnetii* infection. Additionally, by comparing infections with wild-type *C. burnetii* and a *cig2*::Tn mutant having a CCV biogenesis defect that prevents homotypic fusion of lysosome-derived vesicles with the CCV, our data reveal significant differences in pleiomorphism of both the bacteria and CCVs in three dimensions, while demonstrating that proportions of SCV to LCV are nearly identical in cells infected with wild-type and the mutant. Our work demonstrates that FIB-SEM enables the quantitative study of *C. burnetii* infection and opens a new avenue to address various biological questions in diverse intracellular pathogens.

## RESULTS

### FIB-SEM enables quantifiable, three-dimensional visualization of *C. burnetii* infection

We previously used cryo-electron tomography (cryo-ET) to image individual *C. burnetii* cells within the CCV and to observe the development of bacteria over the course of infection (7). However, cryo-ET can only be used to image thin samples, with limited information in the z direction. Here, we used FIB-SEM to obtain a three-dimensional volume of infected HeLa cells and CCVs within, observing a whole population of *C. burnetii*. Immediately apparent are the extremely variable sizes of CCVs formed by wild-type *C. burnetii*, ranging from ∼8 μm^3^ to over 2000 μm^3^ and occupying up to 60% of the imaged volume of the host cell (**Fig. 1**, **2**; **Table 1**). As expected for infection with wild-type *C. burnetii*, five cells in our dataset contained a single CCV. However, one cell appeared to contain two small but separate CCVs (**Fig. 2**). Additional organelles clearly visible in the dataset include mitochondria and nuclei (Fig. **1**). At 12 nm isotropic voxels, our dataset presents a three-dimensional snapshot of *C. burnetii* infection with a singular combination of scale and resolution.

**Figure 1:**
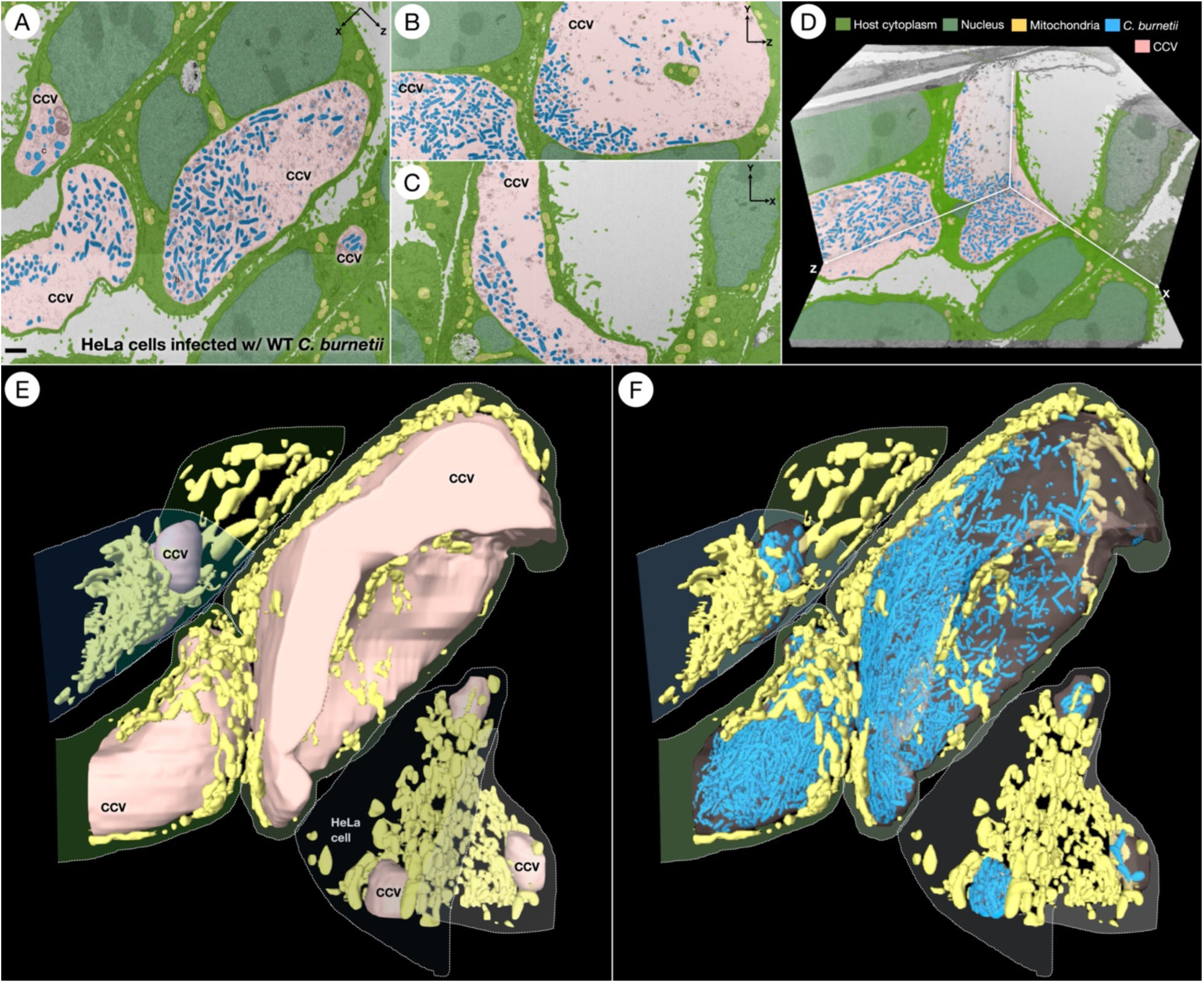
FIB-SEM enables 3D visualization of HeLa cells infected with wild-type *C. burnetii*. (**A-C**) Segmented slices of the FIB-SEM dataset showing HeLa cells infected with wild-type *C. burnetii*. Host cytoplasm (bright green), nuclei (dark green), mitochondria (yellow), *C. burnetii* (blue), CCV (pink). Scale bar 1 μm. (**D**) Three orthogonal slices from the 3D dataset. (**E**) 3D rendering of the CCVs and mitochondria present in the dataset. Individual HeLa cells are outlined with a dashed line. (**F**) The same 3D rendering as shown in panel (**E**). The CCV membranes have been made transparent, and *C. burnetii* cells are visible within CCVs.

**Figure 2:**
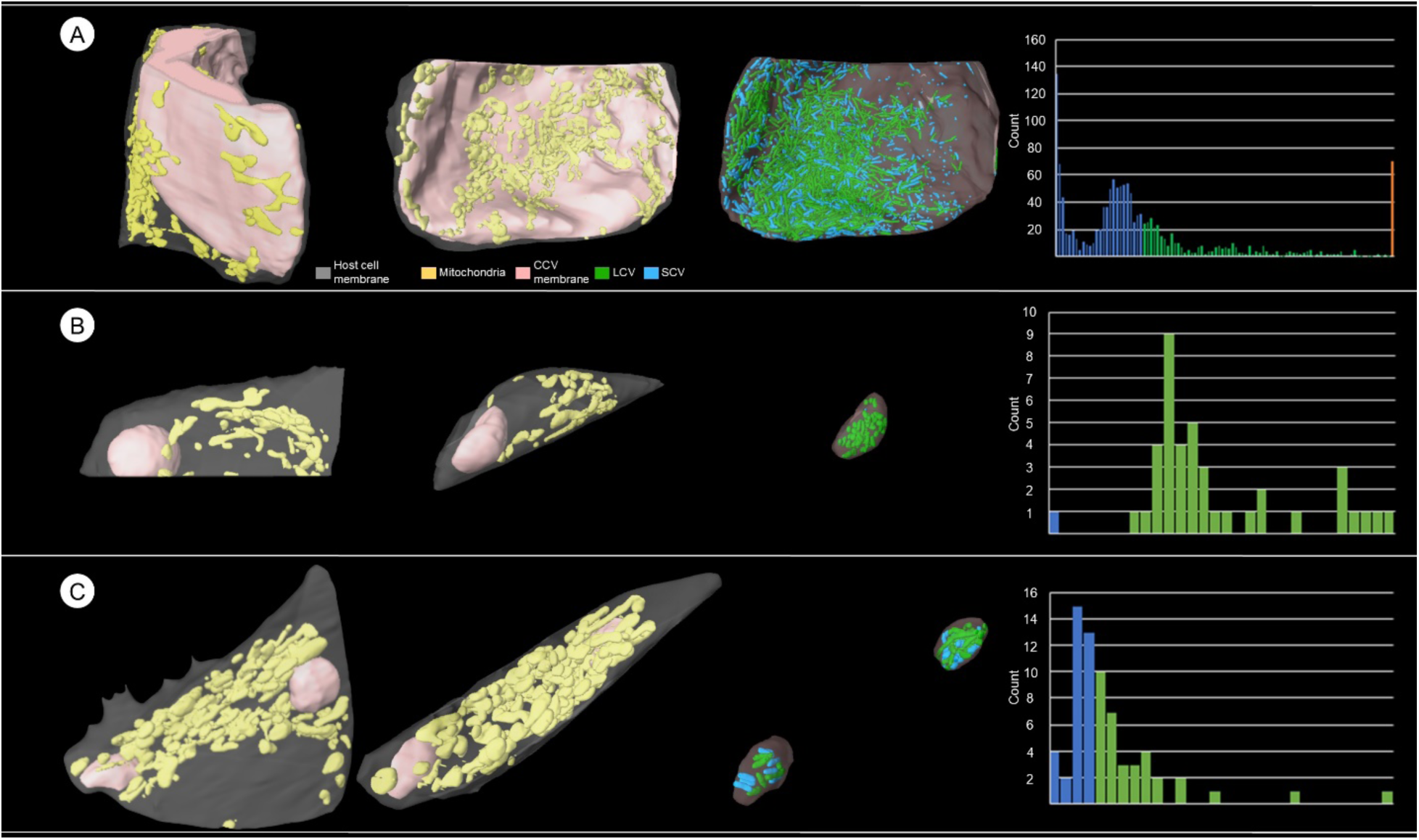
CCV morphology is highly diverse. (**A**) 3D rendering of a HeLa cell containing a large CCV. CCV (pink), mitochondria (yellow), *C. burnetii* SCVs (blue), *C. burnetii* LCVs (green). (**B**) Histogram of *C. burnetii* cell volumes within HeLa cell shown in (**A**). Bars representing SCVs are colored blue, and those representing LCVs are colored green. The largest 5% of cells are grouped together into the orange bar. (**C**) 3D rendering of another HeLa cell with a smaller CCV. (**D**) Histogram of *C. burnetii* cell volumes within HeLa cell shown in (**C**). (**E**) 3D rendering of a HeLa cell containing two small CCVs. (**F**) Histogram of *C. burnetii* cell volumes within HeLa cell shown in (**E**).

**Table 1:** Wild-type statistics.

| CCV | 1 | 2 | 3 | 4 | 5 | 6 | 7 | Average | Total |
| --- | --- | --- | --- | --- | --- | --- | --- | --- | --- |
| CCV volume ($\mu\text{m}^3$) | 8.2 | 19 | 19.2 | 21.8 | 77.2 | 594.7 | 2072.7 | 401.8 | 2812.9 |
| Number of bacteria | 16 | 6 | 27 | 52 | 42 | 961 | 1402 | 358 | 2506 |
| Mean bacterial volume ($\mu\text{m}^3$) | 0.085 | 0.13 | 0.097 | 0.11 | 0.17 | 0.088 | 0.10 | 0.11 | 0.10 |
| Median bacterial volume ($\mu\text{m}^3$) | 0.082 | 0.17 | 0.093 | 0.083 | 0.15 | 0.062 | 0.068 | 0.10 | 0.068 |

### Both *C. burnetii* and CCVs are highly pleiomorphic

To better quantify the characteristics of the bacteria resident in CCVs, we used a convolutional neural network within the Dragonfly software package to segment the entire dataset automatically and accurately, after initial manual segmentation for ground truth generation and training. The resulting model was thoroughly evaluated and then used to identify and characterize 2,506 individual bacterial cells in seven CCVs (**Table 1, Movie 1**).

To characterize the populations of infectious SCVs and replicating LCVs in our dataset, we quantified the volumes of individual cells. The distributions of cell sizes varied greatly among CCVs (**Fig. 2**). These CCVs likely represent a spectrum of different stages of *C. burnetii* infection. A histogram of bacterial cell volumes showed two major peaks, the first at ∼1000 voxels (∼0.007 μm^3^) and the second at ∼8400 voxels (∼0.058 μm^3^). Using a cutoff value of 12,000 voxels (∼0.083 μm^3^) to separate SCVs from LCVs (**Fig. 2**), 1,624 (∼65%) of the cells across all of the CCVs in the dataset were classified as SCVs (**Fig. 3C**). Interestingly, the proportion of SCVs within a vacuole varied greatly among CCVs, from 4.76% up to 70.97%. The two largest CCVs in the dataset contained the largest proportions of small bacteria, 63.98% and 70.97%, consistent with the notion that these larger vacuoles represent later stages of infection (**Fig. 3**).

**Figure 3:**
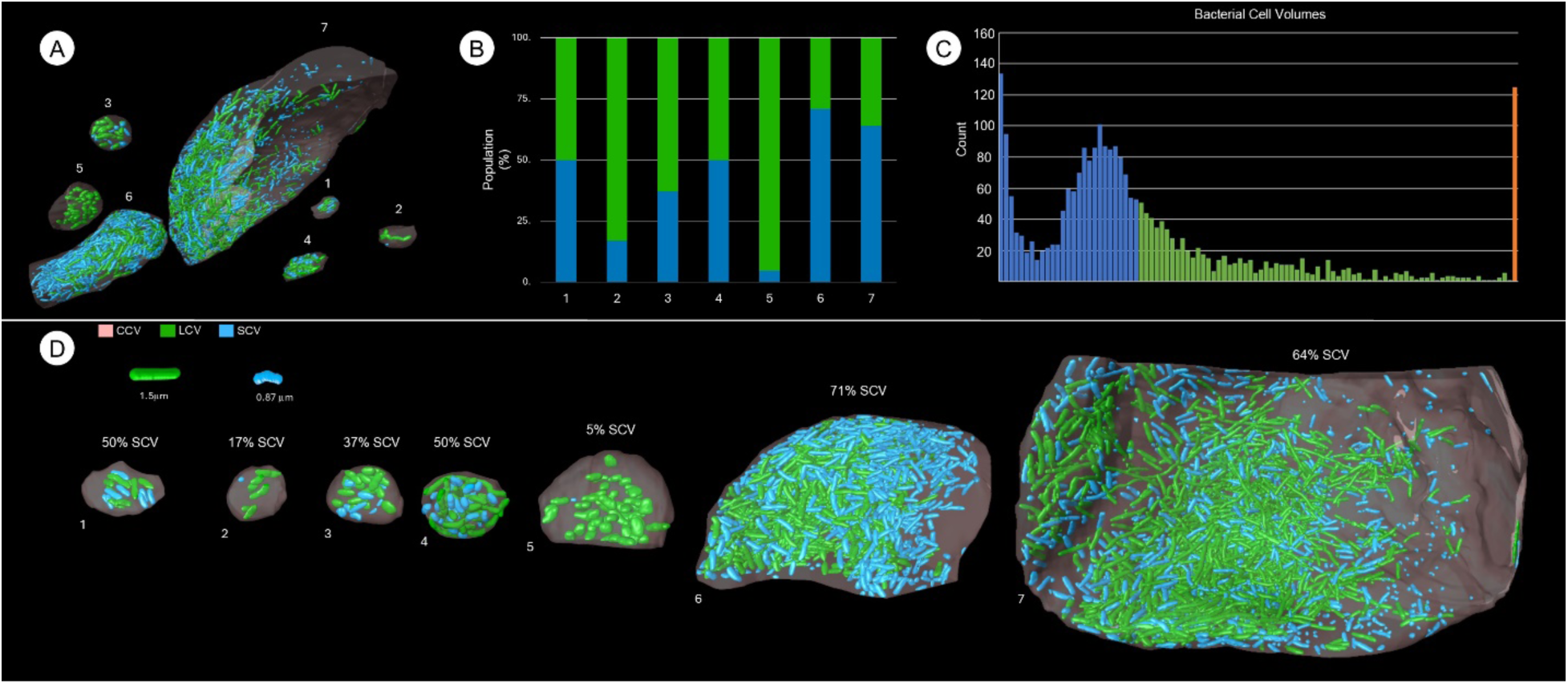
Distribution of bacterial cell volumes varies by CCV. (**A**) 3D rendering of CCVs present in the wild-type FIB-SEM dataset showing the bacteria inside. CCVs are numbered in order by volume. (**B**) Bar graph showing the proportion of SCVs and LCVs in each CCV. (**C**) Histogram of bacterial cell volumes of all seven CCVs. Bars are colored blue (SCV), green (LCV), or orange (>95^th^ percentile). (**D**) Lineup of CCVs showing the bacteria within. Inset: renderings of median LCV and SCV cell. CCV (gray), SCV (blue), LCV (green).

Although cell volumes below the SCV cutoff tended to cluster strongly, volumes above the threshold were much more widely distributed. At the extreme upper end, some clusters of cells are treated as single entities because cells are too close to each other and thus inseparable. To prevent the largest 5% of volumes from distorting the profile, these were binned together in the histogram of the total population’s volumes. Notwithstanding volumes that did not represent individual cells, LCV volumes were still far less consistent than SCV volumes, demonstrating that replicating *C. burnetii* cells are highly pleiomorphic (**Fig. 3**).

Overall, using automated machine learning-based segmentation, we successfully measured several highly variable parameters of bacterial infection, including the range of cell sizes of the notoriously pleiomorphic *C. burnetii*.

### Altered bacterial organization is an observable phenotype of CCV biogenesis defects

Our 3D approach also allowed us to quantify the orientation of each cell relative to a fixed axis. We found that both the phi and theta angles of the *C. burnetii* population are effectively randomly distributed, with no clear preferred orientation (**Fig. 4A-C**). We hypothesized that the cells are free to be buffeted about by microfluidic motion within the large CCVs given that the vacuole contains large amounts of empty space and that, in a more tightly packed CCV, the cells may adopt a single orientation or be otherwise organized differently to increase packing efficiency.

**Figure 4:**
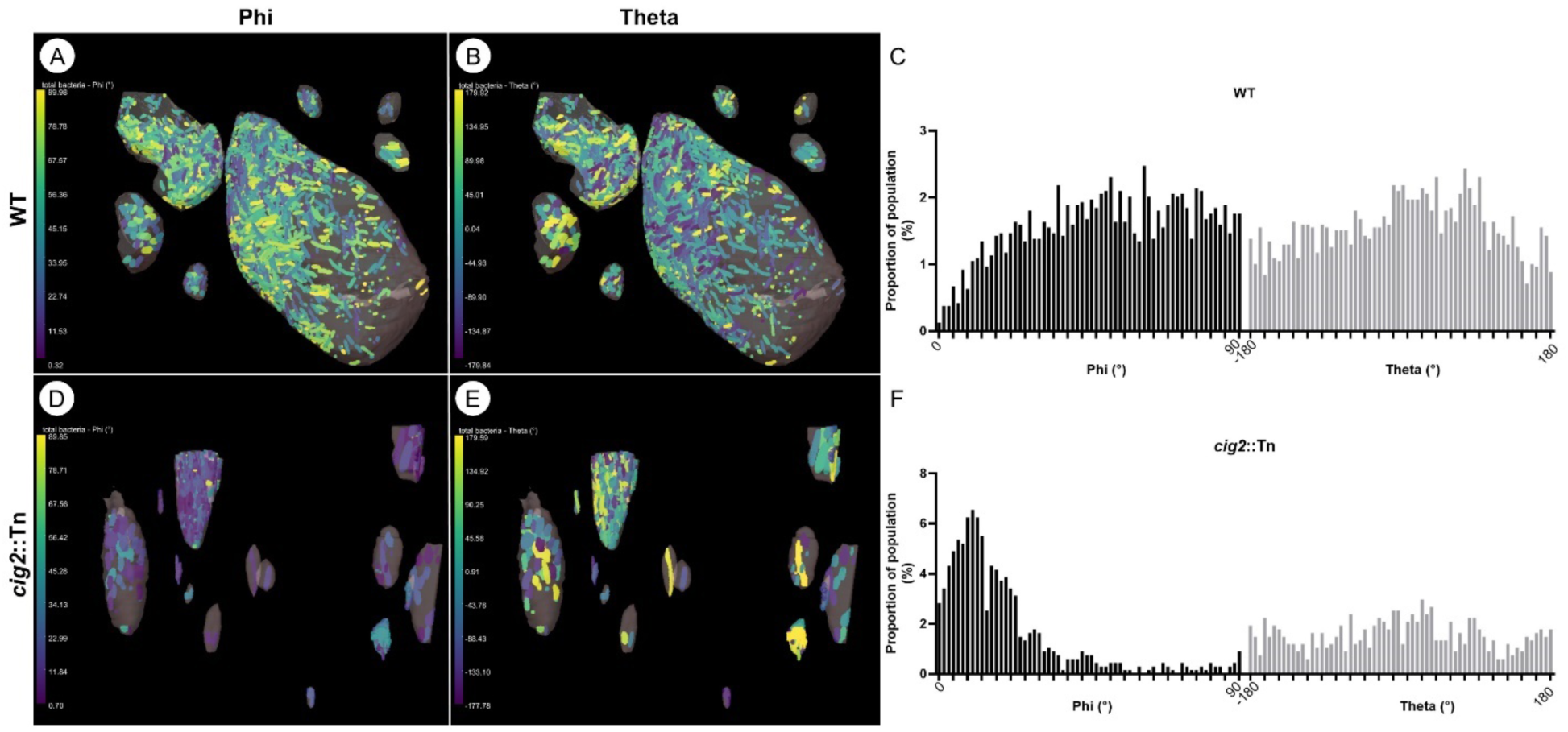
*Cig2*::Tn mutant cells have a preferred orientation within CCVs. 3D representations of wild-type (**A**, **B**) and *cig2*::Tn mutant (**D**, **E**) cells color-coded by their phi (**A**, **D**) or theta (**B**, **E**) angles. (**C**, **F**) Histograms of the phi and theta angles of wild-type (**C**) and *cig2*::Tn mutant (**F**) populations.

To explore this hypothesis, we selected a *cig2*::Tn mutant for direct comparison with wild-type *C. burnetii*. Cig2 is an effector protein promoting constitutive fusion between the CCV and autophagosomes. Cig2-directed biogenesis of an autolysosomal vacuole is essential for the unique fusogenic properties of the CCV and for virulence in an animal model of disease. A *cig2* mutant strain produces smaller CCVs than the wild-type strain and displays a multivacuole phenotype due to a defect in homotypic fusion of individual CCVs within infected cells (14–16). We collected a FIB-SEM dataset from cells infected with a *cig2*::Tn mutant and, in support of our hypothesis, we found that *cig2::*Tn *C. burnetii* cells have a very clear preferred orientation within CCVs, with their phi angles clustered around 11⁰ (**Fig. 4D-F**).

To further characterize bacterial organization, we analyzed the positions of *C. burnetii* cells within the CCV in two ways. First, we counted the number of cells intimately associated with the CCV membrane in each of our datasets. We define “intimately associated” as within 24 nm, the limit of our resolution (**Fig. 5A, B**). We found that the proportion of cells intimately associated with the CCV membrane generally decreased as CCV volume increased, though the correlation was weak (R^2^ ∼0.1) (**Fig. 5C**).

**Figure 5:**
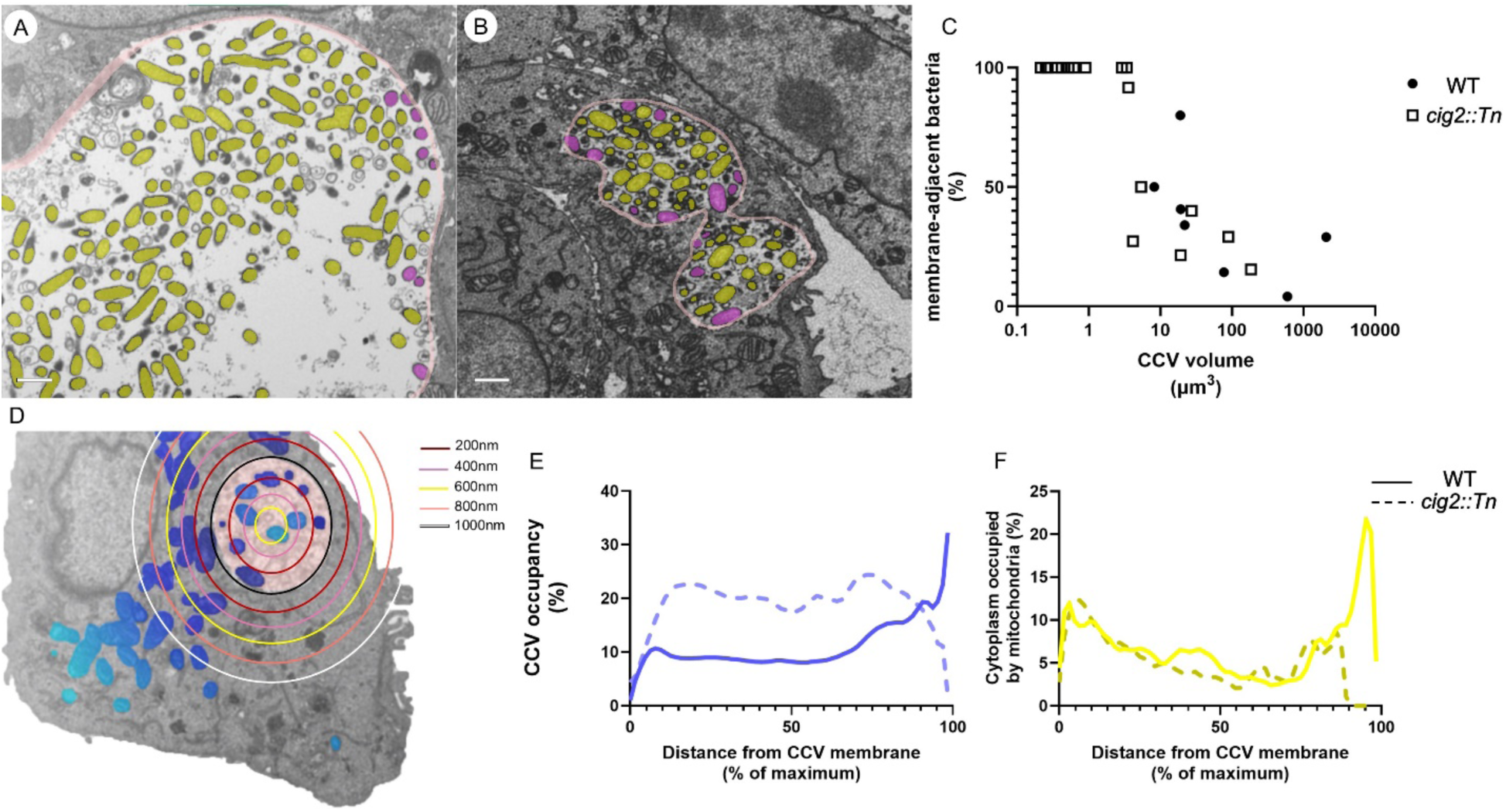
*C. burnetii* shows no tendency to associate with the CCV membrane. Slices of the wild-type (**A**) and *cig2*::Tn mutant (**B**) datasets showing a CCV membrane (pink) and the bacteria inside the CCV. Bacteria have been color-coded depending on whether they are intimately associated with the CCV membrane (purple) or distant from it (yellow). Scale bars 1μm. (**C**) A graph showing the relationship between CCV volume and the percentage of bacteria intimately associated with the CCV limiting membrane. (**D**) A diagram showing a single CCV (pink) and segmented bacteria and mitochondria (blue). Rings are placed at increasing distances from the CCV membrane, and the bacteria and mitochondria are color-coded according to their distance from the membrane. Histograms of the percentage of CCV space taken up by *C. burnetii* (**E**) and cytoplasm space taken up by mitochondria (**F**) binned by distance from the nearest CCV membrane.

Second, we determined the proportion of space taken up by bacteria within the CCV and binned that data by distance from the CCV membrane, creating a picture of the location of bacterial biomass within the CCV (**Fig. 5D-E**). In the region 0.2 μm to 1.8 μm from the membrane in CCVs formed by wild-type *C. burnetii*, bacteria occupy roughly 10% of the CCV volume, but in the area >1.8 μm from the membrane, occupancy rises steadily with distance, up to ∼32% (**Fig. 5E**). Unlike wild-type *C. burnetii*, *cig2*::Tn mutant cells are uniformly distributed throughout the CCVs, taking up roughly 20% of the space (**Fig. 5E**).

In both datasets, we also segmented mitochondria in infected cells to determine if they associate with the CCV during infection, as previously described for instance for intracellular *Salmonella* Typhi (23). As mitochondria are highly interconnected, counting individual mitochondria associated with the CCV is impractical. However, we were able to quantify the percentage of cytoplasmic space taken up by mitochondria as a function of distance from the CCV. For both strains, the distribution of mitochondria in the host cell cytoplasm is very similar (**Fig. 5F**). The percentage of cellular volume occupied by mitochondria as a function of distance from the CCV membrane shows clear peaks in both the CCV-proximal and CCV-distal regions in both the wild-type and mutant datasets (**Fig. 5F**).

Collectively, the organization of the intracellular *C. burnetii* population as measured by preferred orientation and association with the CCV membrane can be readily explained by spatial limitation and thus functions as a quantitative, growth rate-agnostic indicator of CCV biogenesis defects.

### Homotypic fusion is dispensable for the *C. burnetii* developmental cycle

Multiple CCVs were identified within the same host cells infected with the *cig2*::Tn mutant. The CCVs were generally smaller than those formed by wild-type *C. burnetii*, as previously reported (15, 16). Less than 2% of the total host cell volume is comprised by *cig2*::Tn mutant CCVs, much less than the 30% occupied by CCVs in wild-type (**Table 1**, **2**). Some *cig2*::Tn mutant CCVs contain only a single bacterium, and 11/17 contain three or fewer cells (**Fig. 6**, **Table 2**). The smaller CCVs formed by the *cig2*::Tn mutant contain fewer bacterial cells, 689 in total, and the bacteria are smaller than wild-type (**Fig. 6D**, **Table 2**). The *cig2*::Tn mutant volumes below the SCV cutoff form a single peak at ∼2,900 voxels (∼0.02 μm^3^). The population of late-stage, SCV-dominated CCVs in the *cig2*::Tn mutant is also much denser than that of the other *cig2*::Tn mutant CCVs or any CCV in the wild-type sample, with up to 4.3 bacteria/μm^3^ (**Table 2**). Notably, the smallest of these densely packed vacuoles is in close contact with a lysosome, suggesting that bacterial overgrowth may be triggered by signals derived from the lysosome (**Fig. S1**).

**Figure 6:**
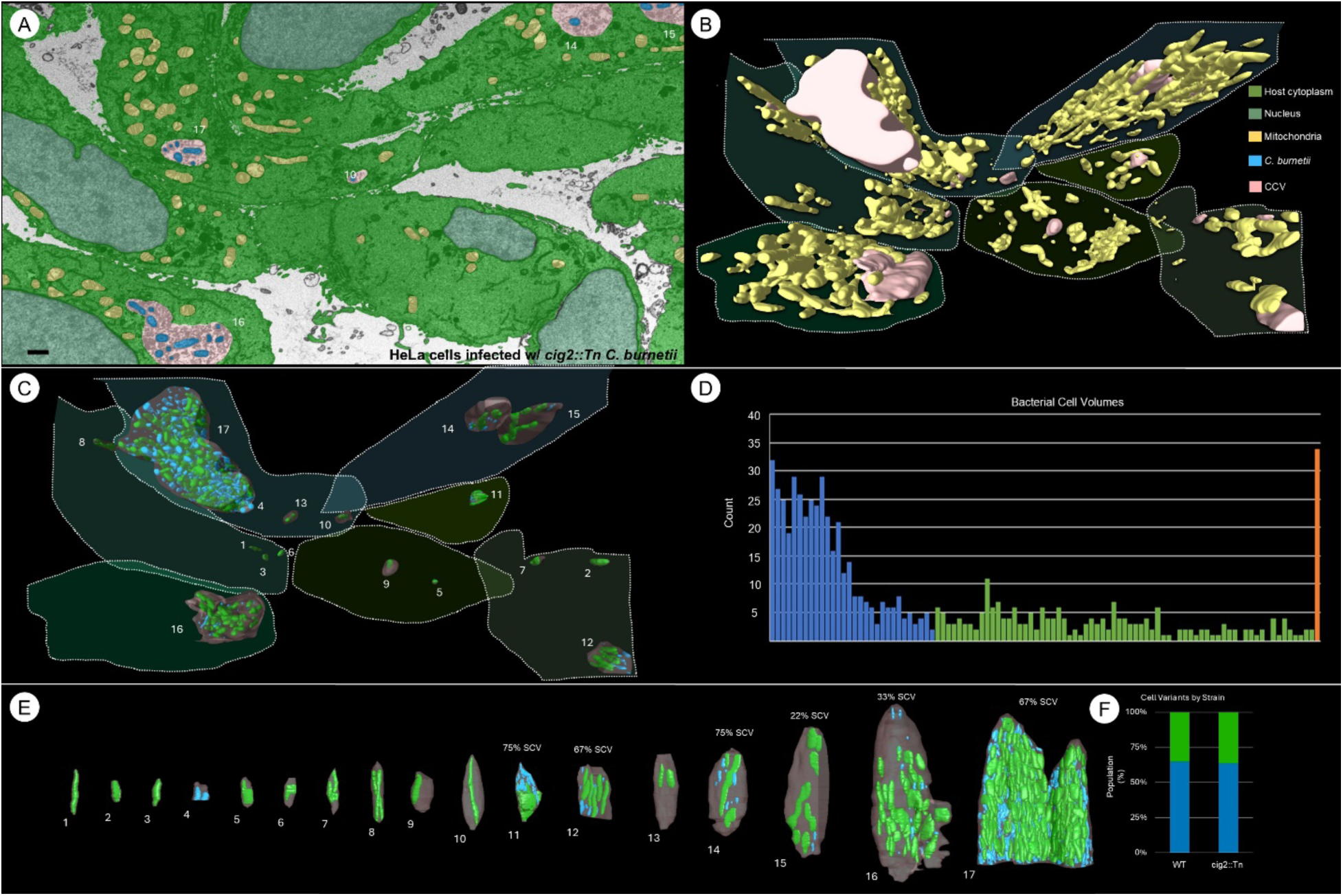
The *cig2*::Tn mutant forms smaller CCVs that contain the same proportions of cell size variants as wild-type. (**A**) A pseudo-colored slice of the *cig2*::Tn mutant FIB-SEM dataset. Host cytoplasm (bright green), nuclei (dark green), mitochondria (yellow), *C. burnetii* (blue), CCV (pink). Scale bar 1 μm. (**B**) 3D rendering of the CCVs and mitochondria present in the dataset. (**C**) 3D rendering of the CCVs present in the dataset and the bacteria inside. CCVs are numbered in order by volume. Individual HeLa cells are outlined. Cells are colored blue (SCV) or green (LCV). (**D**) Histogram of bacterial cell volumes of all 17 CCVs. Bars are colored blue (SCV), green (LCV), or orange (>95^th^ percentile). (**E**) Lineup of CCVs showing the bacteria within. (**F**) Bar graph showing the proportion of SCV and LCV cells in each FIB-SEM dataset.

**Table 2:** *Cig2*::Tn mutant statistics.

| CCV | 1 | 2 | 3 | 4 | 5 | 6 | 7 | 8 | 9 | 10 | 11 | 12 | 13 | 14 | 15 | 16 | 17 | Average | Total |
| --- | --- | --- | --- | --- | --- | --- | --- | --- | --- | --- | --- | --- | --- | --- | --- | --- | --- | --- | --- |
| CCV volume ($\mu\text{m}^3$) | 0.22 | 0.25 | 0.27 | 0.36 | 0.42 | 0.52 | 0.62 | 0.90 | 2.9 | 3.4 | 3.6 | 4.1 | 5.4 | 19.3 | 27.4 | 89.3 | 183.7 | 20.2 | 342.8 |
| Number of bacteria | 1 | 1 | 1 | 3 | 1 | 1 | 2 | 2 | 1 | 1 | 16 | 18 | 2 | 8 | 9 | 52 | 579 | 41 | 698 |
| Mean bacterial volume ($\mu\text{m}^3$) | 0.21 | 0.25 | 0.27 | 0.039 | 0.19 | 0.18 | 0.13 | 0.16 | 0.30 | 0.41 | 0.10 | 0.060 | 0.21 | 0.16 | 0.31 | 0.16 | 0.076 | 0.19 | 0.089 |
| Median bacterial volume ($\mu\text{m}^3$) | 0.21 | 0.25 | 0.27 | 0.033 | 0.19 | 0.18 | | | 0.30 | 0.41 | 0.040 | 0.033 | | 0.042 | 0.25 | 0.14 | 0.041 | 0.16 | 0.045 |

Despite their differences, the wild-type and *cig2*::Tn mutant populations share the same proportions of SCVs and LCVs (**Fig. 6F**). For both strains, these numbers were primarily determined by bacteria in the largest CCV, as these contained more than half of the total bacteria, 56% in the wild-type and 86% in the *cig2*::Tn mutant dataset (**Table 1**, **2**). Surprisingly, regulation of CCV homotypic fusion is not necessary for key parts of the intracellular lifestyle of *C. burnetii*. Overall, our FIB-SEM results provide additional insights into how bacterial subversion of the host autophagy pathway can influence the properties of the CCV and host response to *C. burnetii* infection.

## DISCUSSION

FIB-SEM is emerging as one of the most advanced imaging techniques for the study of infections by intracellular pathogens (24–28). Here, we combined FIB-SEM with machine learning-based data analyses to visualize human cells infected with *C. burnetii* in 3D as well as to yield a far more informative picture of the infection process than previously available at this resolution and scale. Moreover, our newly developed FIB-SEM workflow can be readily applied to diverse infection systems and biological questions.

The CCV is an organelle that matures along the endocytic pathway and is modified by the action of Dot/Icm T4SS effectors to support *C. burnetii* replication (5, 9, 14, 21). One such effector protein, Cig2, promotes fusion of the nascent CCV with autophagosomes (14–16). In the absence of Cig2, infected cells display a multivacuole phenotype (**Fig. 6**) (14–16). Our FIB-SEM data revealed that these vacuoles, along with the bacteria inside them, are much smaller than those formed by wild-type *C. burnetii.* Replication of the *cig2*::Tn mutant seems to outpace CCV expansion, which in turn affects bacterial organization and growth (**Fig. 4**-**6**). We expect that other mutants defective for CCV biogenesis may display similar phenotypes and that analyses of bacterial orientation and localization may be useful tools to dissect the activity of other *C. burnetii* effector proteins.

The quantification of the distribution of both *C. burnetii* cells and host cell mitochondria relative to the CCV membrane revealed an interesting phenotype. In contrast to *Salmonella* infection, for example, during which the *Salmonella*-containing vacuole forms close contacts with host mitochondria (23), with both the bacteria and mitochondria positioned near the vacuole membrane, our data showed that *C. burnetii* primarily localizes away from the CCV membrane. Nevertheless, a subpopulation of mitochondria associates with the CCV, suggesting that *C. burnetii* impacts host cell mitochondrial organization (**Fig. 5F**).

*C. burnetii* displays two distinct morphologies: an infective SCV and a replicative LCV (7, 8, 10). Once an SCV reaches the low pH environment of a late endosome/lysosome, it differentiates into an LCV and begins to divide. In this study, we quantified and visualized, for the first time in three dimensions, the distribution of cell variants across several infected cells. The proportion of SCV to LCV within a given CCV varies widely. While there is no strong correlation between SCV abundance and vacuole size, the two largest vacuoles in the wild-type dataset also had the largest proportions of SCV cells (**Fig. 3**). These vacuoles may represent the final stage of infection. A series of synchronous infection experiments to characterize changes in the population structure and in CCV development over the full course of *C. burnetii* infection would test this hypothesis.

Interestingly, the *cig2*::Tn mutant population did not have a greater proportion of SCVs than wild-type (**Fig. 6F)**. The transition from SCV to LCV is triggered by acidification of the phagosome through fusion with lysosomes during endocytic maturation. This process precedes effector translocation and is thus independent of Cig2 function. However, the signal that causes LCV-to-SCV differentiation is unknown (7). The data presented here suggest that a failure of homotypic fusion neither interferes with nor induces the initial transition from SCV to LCV or the transition from LCV back to SCV. Under a previously proposed model, wherein the LCV-SCV transition is triggered by nutrient depletion (8), our data suggest that nutrient availability is not affected by any phenotype of the *cig2*::Tn mutant, including bacterial density within CCVs. This notion is also consistent with the absence of an intracellular replication defect of this mutant (14, 16).

Overall, this study presents a FIB-SEM workflow that enabled us to visualize large volumes of cells infected by *C. burnetii* and thoroughly characterize the complex lifestyles of the intracellular pathogen *in situ* with unprecedented resolution. We revealed multiple phenomena that warrant further investigation, namely the association of mitochondria with the CCV, association of a densely packed CCV with a lysosome, and variable distribution of cell variants in different CCVs.

## MATERIALS AND METHODS

### Bacterial strains, cell lines, and growth conditions

*C. burnetii* Nine Mile RSA439 (phase II, clone 4) (NMII) (29) and a *cig2* transposon insertion mutant (*cig2*::Tn) (14) were cultured axenically in liquid acidified citrate cysteine medium 2 (ACCM-2; Sunrise Science Products) for 6-8 days at 37°C, 5% CO_2_, and 2.5% O_2_ as previously described (30, 31). When appropriate, chloramphenicol (3 μg/ml) was added to ACCM-2. To calculate multiplicities of infection (MOIs), *C. burnetii* genome equivalents were enumerated by quantitative PCR (qPCR) using *dotA*-specific primers as previously described (14).

HeLa 229 cells (ATCC CCL-2.1) were grown in Dulbecco’s Modified Eagle’s Medium (DMEM) supplemented with 10% heat-inactivated fetal bovine serum (FBS) at 37°C in 5% CO_2_.

### C. burnetii infections

HeLa cells were infected at a high MOI of 250-500. Cells infected with wild-type *C. burnetii* NMII were passaged by trypsinization and diluted 1:3 approximately every 3 days for about 2 months prior to fixation for microscopy. Cells infected with the *cig2*::Tn mutant were first expanded by trypsinization and then diluted 1:2 once prior to fixation for microscopy on day 8 post-infection.

### Focused ion beam-scanning electron microscopy (FIB-SEM)

Infected cells were fixed in 2.5% glutaraldehyde and 2% paraformaldehyde in 0.1 M sodium cacodylate buffer pH 7.4 containing 2% sucrose for 1 hour, then rinsed in buffer, and replaced with 0.1% tannic acid in buffer for another hour. Samples were further postfixed in 1% osmium tetroxide and 1.5% potassium ferrocyanide in buffer for 1 hour. The samples were rinsed in sodium cacodylate and distilled water and en bloc stained in 0.5% aqueous uranyl acetate overnight, followed by further rinsing in distilled water and then placed in 60 °C lead aspartate for 1 hour. After 1-hour rinsing in distilled water, the samples were dehydrated through an ethanol series to 100% and were infiltrated with Embed 812 (Electron Microscopy Sciences) resin, placed in silicone molds, and baked at 60°C for at least 24 hours.

The resin block was trimmed to rough area of interest and the surface was cleanly cut. The entire pyramid was carefully removed with a fine blade and mounted on an aluminum stub using conductive carbon adhesive and silver paint (Electron Microscopy Sciences) and then sputtered with approximately 20 nm Pt/Pd (80/20) using a Cressington HR sputter coating equipment (Ted Pella, Inc., Redding, CA, USA) to reduce charging effects.

A dual-beam focused ion beam-scanning electron microscope (FIB-SEM) (ZEISS Crossbeam 550) with a Gallium ion source was used to mill the samples, and a secondary electron detector was used for imaging. SmartSEM (ZEISS, Jenna Germany) was used to set up initial parameters and to find the regions of interest (ROI) by SEM images at 10 kV. 50 μm width X 30 μm height X actual depth was 30 μm with 7 nm/pixel and 7 nm per slice. A platinum protective pad was deposited at the ROI with the FIB (30 kV, 50 pA) to protect the structure and reduce charging. Carbon deposits mill, and highlight was done at 30 kV and 3 nA. A coarse trench milled 30 kv 30 nA followed by fine milling at 30 kV 3 nA. For final acquisition of a cuboid, the area of interest was milled at 30 kV and 300 pA. After milling each slice, an image was acquired by a backscattered electron detector (acceleration voltage of 1.5 kV, imaging current of 2 nA, and aperture diameter of 100 μm) with a pixel dwell time of 2 μs. Atlas 5 (ZEISS) was used for preliminary image stack alignment, and the image data was exported in Tiff format.

### Segmentation, visualization, and data analysis

The FIB-SEM images were imported and analyzed using Dragonfly software (Version 2022.2, Comet Technologies Canada Inc., Montreal, Canada) (32). Automatic data segmentation was achieved using convolutional neural networks with U-net architecture. Ground truth was generated by manual segmentation of 5-10 slices before models were trained. Models trained to segment bacteria were applied only to volumes annotated as CCV, while models to segment mitochondria were applied to whole datasets. It was necessary to train separate models for each dataset. Minimal postprocessing was used to correct automatic segmentations. Volumes and positions of annotated entities were quantified using the connected components analysis feature within Dragonfly. Dragonfly was also used to generate all images and movies of segmented data.

## ACKNOWLEDGMENTS

This work was funded by the National Institutes of Health (NIH) grant R01AI152421 to Jun Liu and Craig R. Roy. The Zeiss Crossbeam 550 FIB-SEM system was acquired through a Major Instrument Grant from NSF #1725480 to Pietro De Camilli, Derek Toomre, and Xinran Liu. We thank Jennifer Aronson for critical reading of the manuscript.

---

**Figure S1:**
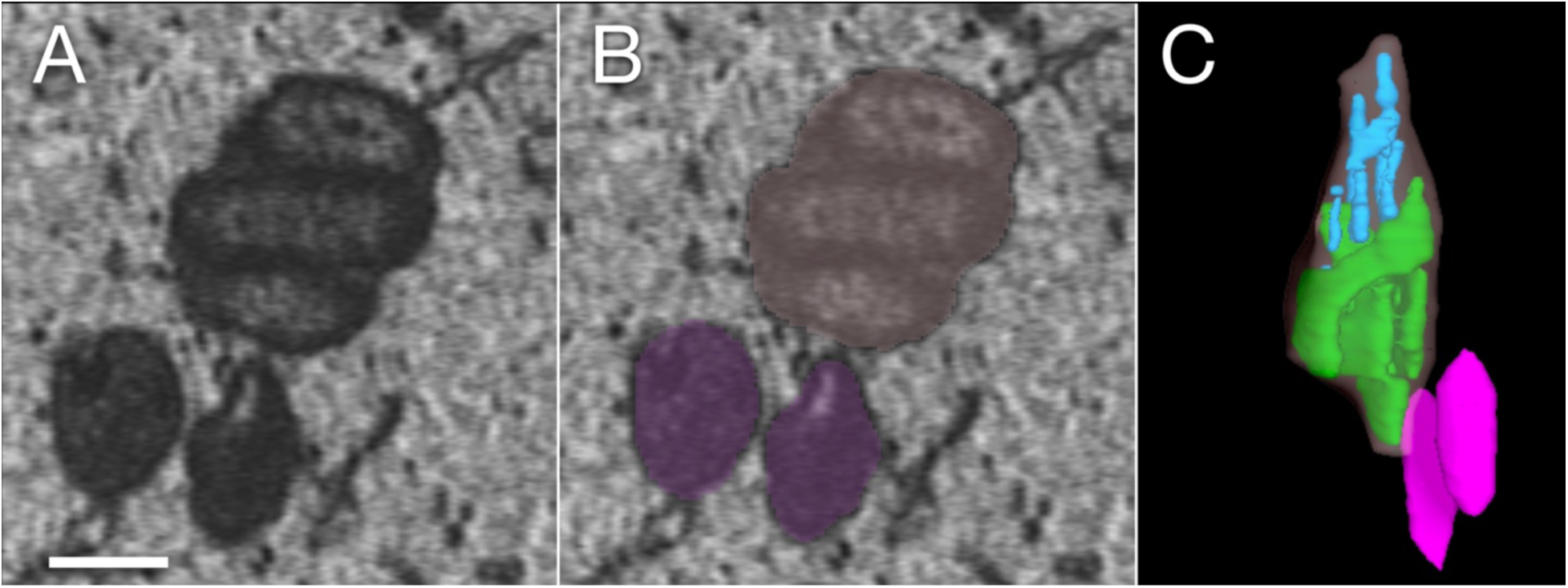
A CCV adjacent to lysosomes. **(A)** A slice from the *cig2*::Tn mutant dataset showing a CCV in contact with a lysosome. **(B)** CCV (gray) and lysosomes (purple) are segmented, respectively. **(C)** 3D rendering of the CCV (gray) and lysosomes (purple). There are several *C. burnetii* cells (green, blue) in the CCV. Scale bar 1μm.

